# Cooperative Non-bonded Forces Control Membrane Binding of the pH-Low Insertion Peptide pHLIP

**DOI:** 10.1101/341628

**Authors:** C. Gupta, Y. Ren, B. Mertz

## Abstract

Peptides with the ability to bind and insert into the cell membrane have immense potential in biomedical applications. pH (Low) Insertion Peptide (pHLIP), a water-soluble polypeptide derived from helix C of bacteriorhodopsin, can insert into a membrane at acidic pH to form a stable transmembrane *α*-helix. The insertion process takes place in three stages: pHLIP is unstructured and soluble in water at neutral pH (state I), unstructured and bound to the surface of a membrane at neutral pH (state II), and inserted into the membrane as an *α*-helix at low pH (state III). Using molecular dynamics (MD) simulations, we have modeled state II of pHLIP and a fast-folding variant of pHLIP, in which each peptide is bound to a 1-palmitoyl-2-oleoyl-*sn*-glycero-3-phosphocholine (POPC) bilayer surface. Our results provide strong support for recently published spectroscopic studies, namely that pHLIP preferentially binds to the bilayer surface as a function of location of anionic amino acids and that backbone dehydration occurs upon binding. Unexpectedly, we also observed several instances of segments of pHLIP folding into a stable helical turn. Our results provide a molecular level of detail that is essential to providing new insights into pHLIP function and to facilitate design of variants with improved cell-penetrating capabilities.

## INTRODUCTION

Oncology is difficult from the perspective of both diagnosis and treatment: early detection is often impossible to achieve (e.g., pancreatic cancer) (1) and conventional treatments suffer from off-target side effects in which chemotherapeutics attack healthy as well as cancerous cells (2). New approaches to treatment aim to increase the effectiveness of targeting, as evidenced by monoclonal antibodies (3) and antibody drug conjugates (ADCs) (4), which distinguish between cancerous and healthy cells by binding cancer-specific antigens. However, this approach has potential shortcomings, due to heterogeneity of tumor cells and rapid mutations of targeted antigens, leading to resistance. An alternative targeting method is to utilize unique characteristics of the tumor microenvironment for exploitation (e.g., hypoxia or acidity in the exterior of the cell). The extracellular environment of tumor cells (pH 6.5-7.0) is more acidic than normal tissue (pH 7.2-7.5) (5). Tumor acidosis stems from altered and enhanced metabolism within the cancer cell. This is manifested through the tendency of cancer cells to utilize glycolysis for ATP synthesis (i.e., the Warburg effect (6)), leading to an excess of lactic acid that is pumped outside the cell. One facet of acidosis that makes it an appealing targeting factor is the fact that it is a universal feature of all cancer cells. Developing a therapeutic approach targeting this property would potentially have far-reaching effects in oncology.

One approach is the pH (Low) insertion peptide (pHLIP) (7, 8). pHLIP resides in three distinct states that are dependent upon the surrounding environment: unstructured and soluble in water at alkaline pH (state I); unstructured and bound to the cell membrane surface at alkaline pH (state II); and inserted across the membrane as an *α*-helix at acidic pH (state III) (Fig. 1). Peptide insertion is induced by a shift from alkaline to acidic pH, with an apparent pK of insertion in 1-palmitoyl-2-oleoyl-snglycero-3-phosphocholine (POPC) membranes of ~6.1 (9). This shift in pH spurs the protonation of two acidic residues (D14 and D25) in the membrane-spanning region of the peptide, leading to an increase in hydrophobicity and triggering the folding and insertion of the peptide across the lipid bilayer (9, 10). The presence of two tryptophan residues (W9 and W15) allow intrinsic fluorescence spectroscopy to be used to detect the transition of pHLIP from state I → II → III.

**Figure 1:**
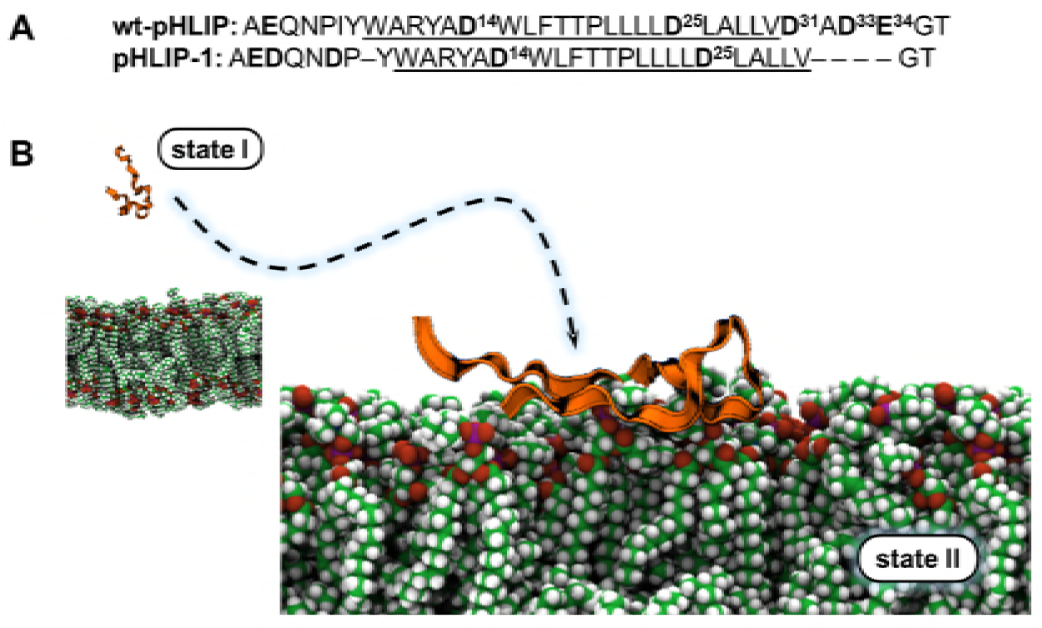
Systems investigated in this study. A) Primary sequence of wt-pHLIP and pHLIP-1. *Boldface*: acidic residues in the sequence; *underline*: putative region that undergoes folding into a transmembrane helix. B) Schematic of how pHLIP binds to a plasma membrane surface. As pHLIP diffuses in solution, it can take on many conformations (state I, *left*). pHLIP will then encounter the membrane surface and partition into the headgroup region (state II, *right*). *Ribbons*: pHLIP; *spheres*: lipids in a POPC bilayer.

Our understanding of the state I → state II transition – binding of pHLIP to the membrane surface – has evolved since pHLIP was first discovered, due to advances in the techniques used to characterize pHLIP as well as the lipid systems being studied. Isothermal titration calorimetry experiments on pHLIP and 1-palmitoyl-2-oleoyl-*sn*-glycero-3-phosphocholine (POPC) vesicles determined that the most important thermodynamic step in pHLIP function is binding to the membrane surface (AG_bind_ = −7.2 kcal-mol^−1^, whereas ΔG_insert_ = −1.8 kcal-mol^−1^) (9). This binding is accompanied by burial of W9 and W15 due to desolvation of pHLIP, a necessary prerequisite for the state II → state III transition (11, 12). Solid-state NMR studies have also shown that 1) residue-specific differences in pHLIP-membrane interactions occur at each of three pH values (pH 7.4, 6.4, and 5.3) (13) and 2) each aspartic acid residue in pHLIP has a distinct p*K_A_*, with the C-terminal residues being titrated first (14). When adding anionic phospholipids (i.e., phosphatidylserine) to the vesicles, the free energy of binding is essentially the same (15). However, Ladokhin and coworkers determined that state II is spectroscopically silent in the presence of non-PC lipids – pHLIP will bind to lipid vesicles but does not partition below the headgroup region, leading to a lack of change in tryptophan fluorescence from state I to state II (16, 17). These more recent studies strongly suggest that a more complex mechanism of binding exists for pHLIP. Despite these efforts to characterize state II, knowledge of the detailed interactions between pHLIP and the cell membrane that eventually lead to folding and insertion are sorely lacking. Without this knowledge, it will be extremely difficult to intelligently design pHLIP variants with the ability to modulate apparent p*K*’s of insertion to closely match the pH of acidic microenvironments such as cancer cells.

Molecular dynamics (MD) simulations are capable of providing atomistic details on the biophysical interactions between pHLIP and a lipid bilayer that are unattainable through experimental methods. In this study we have carried out equilibrium MD simulations to characterize the biophysical interactions that govern the behavior of pHLIP (wt-pHLIP) and a fast-folding variant of pHLIP (pHLIP-1) in state II. Our goal was to address the following unresolved issues with respect to state II: 1) Is there a favored binding complex? 2) What is the mechanism by which this binding occurs? 3) Does partitioning-folding coupling occur during binding? 4) If so, is folding conserved in a specific region of pHLIP? We chose to study wt-pHLIP and pHLIP-1 (18) in order to compare the canonical version of pHLIP with a variant that possesses helical character in state II. Our simulations capture the non-bonded interactions that lead to binding of both peptides, showing good agreement with recent experiments and identifying common motifs in the binding process among different orientations of pHLIP with respect to the bilayer surface. We observe stable formation of *α*-helical folds in pHLIP-1 centered around the D14 and D25 protonation switches; surprisingly, wt-pHLIP also shows an ability to adopt stable helical turns. Most importantly, a clear difference exists in the effectiveness of binding of wt-pHLIP and pHLIP-1 to POPC, indicating that peptide orientation has a direct effect on promoting formation of pHLIP-bilayer complexes.

## MATERIALS AND METHODS

Previously we carried out MD simulations of wt-pHLIP in implicit solvent to determine the most likely conformation of pHLIP in state I (19). We selected a representative snapshot from our most populated cluster in our K-means clustering analysis (i.e., with the lowest root-mean squared deviation to the average structure in the cluster) as the starting structure of wt-pHLIP for our state II simulations. pHLIP-1 (based on the sequence from Karabadzhak et al. (18)) was modified from helix C of the crystal structure of bacteriorhodopsin (PDB ID: 2NTU). To drive pHLIP-1 into a coiled conformation, an MD simulation in the *NVT* ensemble (*T* = 700K) was conducted *in vacuo* for 2 ns using a 2.0 fs timestep using NAMD 2.9 (20). This structure was then solvated in implicit solvent as in (19) and simulated for 2 *µ*s. K-means clustering was then used to select the most probable conformation to form the peptide-bilayer complex.

A POPC bilayer of 200 lipids was prepared using the membrane builder in CHARMM-GUI (21–26) and equilibrated in the *NPT* ensemble (*T* = 310 K, *P* = 1 atm) for 50 ns using NAMD 2.10 and the c36 lipid force field (27). A 2.0 fs timestep, with a force-based switching function for Lennard-Jones interactions from 10 to 12 Å, Langevin thermostat, Nosé-Hoover barostat, and a flexible cell with constant ratio were used. wt-pHLIP and pHLIP-1 structures were then merged with the equilibrated POPC bilayer to generate five independent peptide-bilayer complexes for each type of peptide, referred to as 0°, 72°, 144°, 216° and 288°, with degrees measured as the rotation of each respective peptide around their first principal component axis. Note that these angles are arbitrary and do not correlate between wt-pHLIP and pHLIP-1 simulations. Overlapping waters and lipids were removed. The final systems had a lipid:peptide ratio ~200:1 (exact ratio varied depending on how many lipids had to be eliminated for each system), with a slight degree of bilayer asymmetry.

Unbiased molecular dynamics simulations in the tensionless *NPT* ensemble were performed with the GPU-accelerated version of pmemd in Amber 16 (28, 29) using the CHARMM 36 force field including a modification to better account for cation-pi interactions (27, 30, 31). A hard cutoff of 8 Å was applied to non-bonded forces, as recommended by the CHARMM community (23). The particle mesh Ewald (PME) approach was used for computing electrostatic forces (32, 33), and hydrogens were restrained with the SHAKE algorithm (34). Simulations were run with a 2.0 fs timestep at 310 K with a Langevin thermostat. The Berendsen barostat was used to maintain pressure at 1 atm with semi-isotropic pressure scaling. Visualization was performed using VMD (35), data analysis was carried out using VMD and in-house python scripts, and matplotlib and gnuplot were used for making plots.

## RESULTS

### Partitioning of pHLIP influenced by location of acidic residues

The recent NMR work of An and Qiang examined partitioning of pHLIP into POPC vesicles during the state II → state III transition over incremental pH jumps (pH 7.4 → 6.4 → 5.3) using rotational-echo double-resonance (REDOR) NMR spectroscopy (13). The C-terminal residues that were isotopically labeled (L21, L22, L26, and A27) displayed a high degree of dynamics, leading to their hypothesis that the polar C-terminus of pHLIP (D31, D33, and E34) plays a major role in preventing partitioning of that end of pHLIP into the bilayer surface. A subsequent follow-up study from An and Qiang showed that the acidic residues in the C-terminus of pHLIP not only affect the ability for pHLIP to partition into the bilayer surface, but they are also the first residues to be titrated upon acidification of the surrounding environment (14). What was unclear was the specific relationship between binding of pHLIP to the bilayer surface and titration of acidic residues – is titration facilitated by a change in hydrophobicity of the surrounding environment through burial in the headgroup region, or does exposure to solvent, as most likely with the C-terminus of pHLIP, allow for faster transitions from the acidic to the neutral state? Our simulations on wt-pHLIP and pHLIP-1 consistently remain bound to the bilayer surface, but with significant distinctions between the two peptides and between orientations. For wt-pHLIP, the 0° and 72° orientations utilize a combination of salt bridge formation with R11 and partitioning of aromatic sidechains within the N-terminal segment of the peptide to effectively bind to the bilayer surface (Fig. 2A). However, the 144° orientation completely dissociates from the bilayer around 300 ns, while the 216° and 288° orientations establish stable binding via the sinker stretch of nonpolar residues from position 21-30. In addition, the hydrophobic sinker stretch from T18 to L24 is able to remain embedded within the bilayer for all of the bound orientations of wt-pHLIP, indicating that it plays a significant role in the quick cooperative response in state II that was observed by An and Qiang when shifting from pH 7.4 to pH 6.4. Unlike wt-pHLIP, pHLIP-1 remains stably bound to the bilayer surface regardless of orientation. Binding is dominant from position 10 to the C-terminus and is a significant extension of the binding region of the sinker stretch (Fig. 2B). The most likely explanation for this stability in binding is due to the lack of acidic residues on the C-terminus of pHLIP-1.

**Figure 2:**
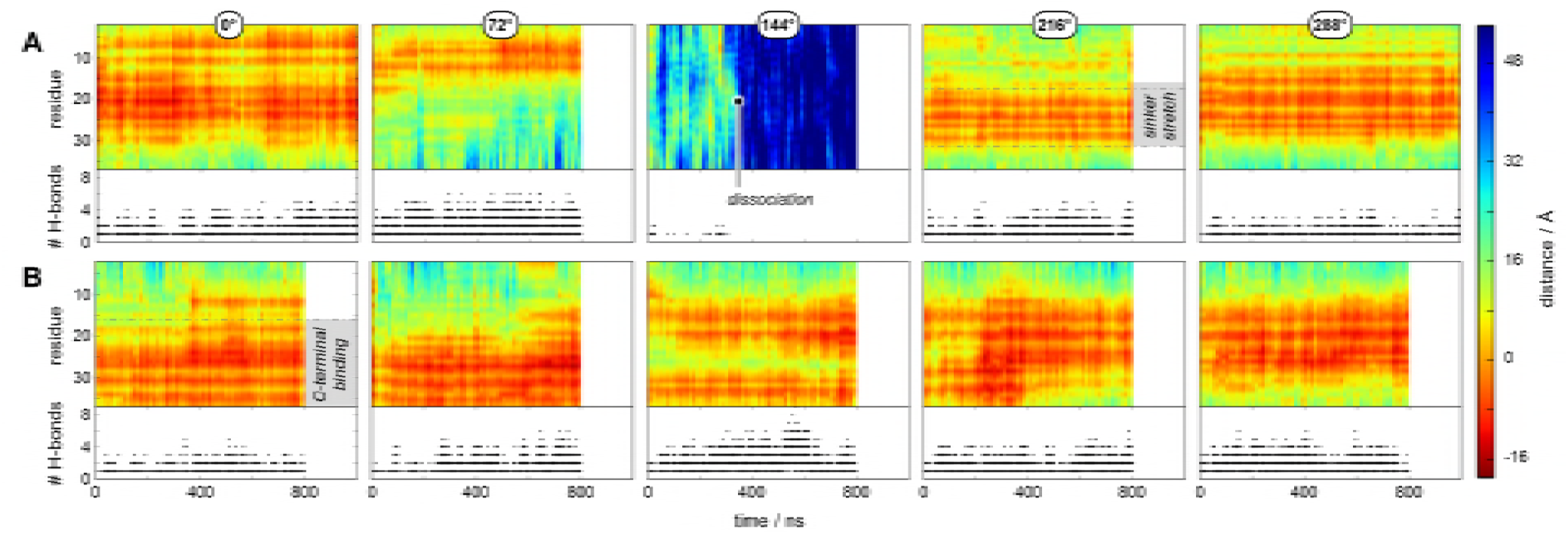
Quality of binding of pHLIP can depend on orientation and peptide composition. A) Per-residue distance of wt-pHLIP from the center of the POPC bilayer as a function of time. wt-pHLIP can bind to the surface of a POPC bilayer through electrostatic or non-polar interactions. In the case of 0° and 72° orientations, the N-terminal portion of pHLIP binds stably (over hundreds of ns), mainly due to a combination of salt bridge formation with R11 and partitioning of aromatic sidechains (tyrosine and tryptophan). The 144° orientation is transiently bound to the bilayer surface and completely dissociates around 300 ns. In contrast, the 216° and 288° orientations remain stably bound to the bilayer surface, this time through partitioning of the nonpolar “sinker stretch” of the C-terminal half of the peptide. B) Per-residue distance of pHLIP-1 from the center of the POPC bilayer as a function of time. In general, the opposite behavior is observed for pHLIP-1. C-terminal residues predominantly bind to the POPC surface for extended periods of time (hundreds of ns). For the most part, binding of pHLIP-1 is independent of orientation; see Fig. S1 for details on the distribution of binding.

Our results qualitatively agree with the aforementioned NMR studies; each respective half of the peptide with terminal polar residues (N-terminus for pHLIP-1, C-terminus for wt-pHLIP) lies furthest from the bilayer surface. If we group wt-pHLIP and pHLIP-1 by region (N-terminus: residues < 10; middle: residues 10-27; C-terminus: residues > 27), more general patterns begin to emerge. The middle segment in wt-pHLIP most consistently binds to POPC. In addition, the N- and C-terminal segments of wt-pHLIP have opposite binding affinities to POPC: the N-terminus, with only one charged residue, binds more effectively, while the C-terminus, with three acidic residues, binds poorly or not at all (Fig. S1A, Supporting Information). Unlike wt-pHLIP, binding of pHLIP-1 is fairly consistent between all five orientations for both the N-terminal and middle segments — no binding of the N-terminus (higher concentration of acidic residues) and effective binding of the middle segment. For the C-terminal segment of pHLIP-1, binding also occurred in the majority of the simulations, with a larger distribution (Fig. S1B, Supporting Information). Each orientation of the C-terminus has equivalent or improved binding effectiveness in pHLIP-1 compared to wt-pHLIP. The solid-state NMR study of An and coworkers determined the distances of isotopically-labeled alanine residues (A10, A13, and A27) in wt-pHLIP from the upper leaflet of POPC vesicles in order to characterize the quality of binding of these residues in state II. In most cases of our wt-pHLIP simulations, A10 and A13 do not fully reproduce these equilibrium distances (Fig. S2A, Supporting Information). However, it is of note that the lipid to peptide ratio used in the NMR experiments was 75:1, whereas our simulations were conducted at a ratio of 200:1. Previous studies have shown that binding of pHLIP is affected by lower lipid:peptide ratios, leading to a “parking problem” for pHLIP on the bilayer surface (9). For the orientations that remained bound to POPC, the 0°, 72°, and 288° orientations all had distance distributions for A27 that were > 10 Å in agreement with the NMR experiments. For pHLIP-1, distances are much closer to the POPC surface than wt-pHLIP for all three alanine residues (Fig. S2B, Supporting Information). This is in part because A27 is not proximal to the three C-terminal acidic residues present in wt-pHLIP, which has the potential for electrostatic repulsion with the negatively-charged phosphate region of the PC headgroups. In addition, the distances for A10 and A13 are slightly reversed; A13 is closer to the P atoms than A10. This reversal could be due in part to the presence of negatively-charged residues at positions 2, 3, and 6 in pHLIP-1, which could have a more marked repulsive effect on the closer alanine residue (i.e., A10).

### pHLIP samples secondary structural conformations in state II

The conventional model of state II proposes that the peptide remains in a coiled conformation when bound to a lipid bilayer at neutral or alkaline pH. For our simulations of wt-pHLIP, this is predominantly the case. However, we also observed that wt-pHLIP can form stable *α*-helices for hundreds of ns (Fig. 3A). In particular, the 0° and 288° orientations of wt-pHLIP undergo helical folding in multiple regions of the N-terminal half of the peptide, with turns centered around W9 and F17. There is precedence for this phenomenon, as deep-UV resonance Raman (DUVRR) spectroscopy experiments showed that pHLIP can adopt helical structure in state II (Cooley and Jiji BC 2014). In contrast, pHLIP-1 undergoes helical folding regardless of orientation (Fig. 3B). This is consistent with CD spectra that showed pHLIP-1 possesses a noticeable degree of helicity in state II (Karabadzhak BJ 2012). Folding occurs in an extended segment from R11 to T19, and for three of the orientations (0°, 72°, and 216°), the C-terminus folds into a helix. In general, any occurrence of secondary structure is helical in nature (Supporting Information Fig. S3). Visual inspection of our simulations shows that for both wt-pHLIP and pHLIP-1, partitioning via sidechain-headgroup interactions is most prevalent with the more non-polar residues (i.e., leucine and isoleucine). The role of aromatic and charged sidechains in binding of pHLIP to a POPC bilayer surface can be noticeably different between the two peptides, as will be discussed below.

**Figure 3:**
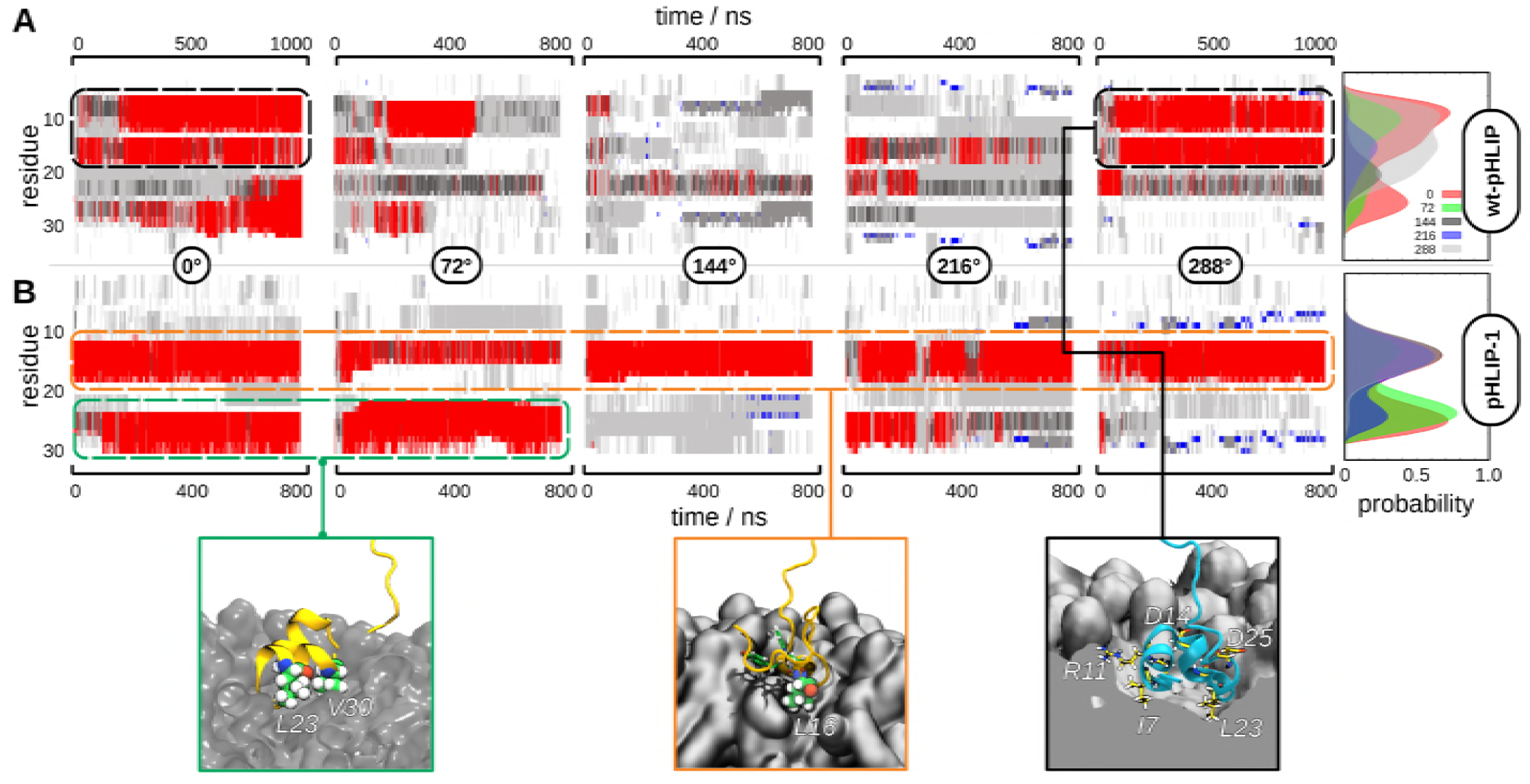
Formation of helical turns occurs during binding for both wt-pHLIP and pHLIP-1. **A)** *Left*: time-dependent secondary structure calculation of the different orientations of wt-pHLIP (0°, 72°, 144°, 216°, and 288°) when bound to POPC. wt-pHLIP can form stable *α*-helices for hundreds of ns. In particular, the 0° and 288° orientations of wt-pHLIP undergo helical folding in multiple regions of the N-terminal half of the peptide (*black dashed line*). *Right*: per-residue distribution of *α*-helical folding of wt-pHLIP when bound to POPC. A residue is defined to be forming an *α*-helix if it is a part of a helical stretch of 3 or more residues. *Red*: *α*-helix; *blue*: *β*-sheet; *gray*: coiled conformation. **B)** *Left*: time-dependent secondary structure calculation of the different orientations of pHLIP-1 (0°, 72°, 144°, 216°, and 288°) when bound to POPC. The propensity of pHLIP-1 to fold into an *α*-helix is even more pronounced than for wt-pHLIP: the middle segment of pHLIP-1 undergoes stable formation of an α-helical turn in all orientations (*orange dashed line*). In addition, in two of the five orientations, the C-terminal segment of pHLIP-1 undergoes folding into an α-helix (*green dashed line*). *Right*: per-residue distribution of *α*-helical folding of pHLIP-1 when bound to POPC. Orientation influences the relative distribution of helix-formation between the two putative helix-forming domains but the overall pattern is invariant to orientation. Color scheme is the same as in A.

This stark contrast in the onset of folding in state II is due to several factors. One of the most significant that we observed is formation of an *i*+3 salt bridge between R11 and D14 in pHLIP-1. All simulations of pHLIP-1 show stable formation of this salt bridge with little to no fluctuations over hundreds of ns (Fig. 4). The R11-D14 interaction also directly coincides with the region of pHLIP-1 that folds into an *α*-helix. In the wt-pHLIP simulations, fewer salt bridges are formed and are very transient; no *i*+3 salt bridges occur (Fig. 4). There appears to be no dominant intramolecular interaction, as R11 in wt-pHLIP interacts transiently with all of the negatively-charged residues, depending on the orientation of the peptide in the bound complex.

**Figure 4:**
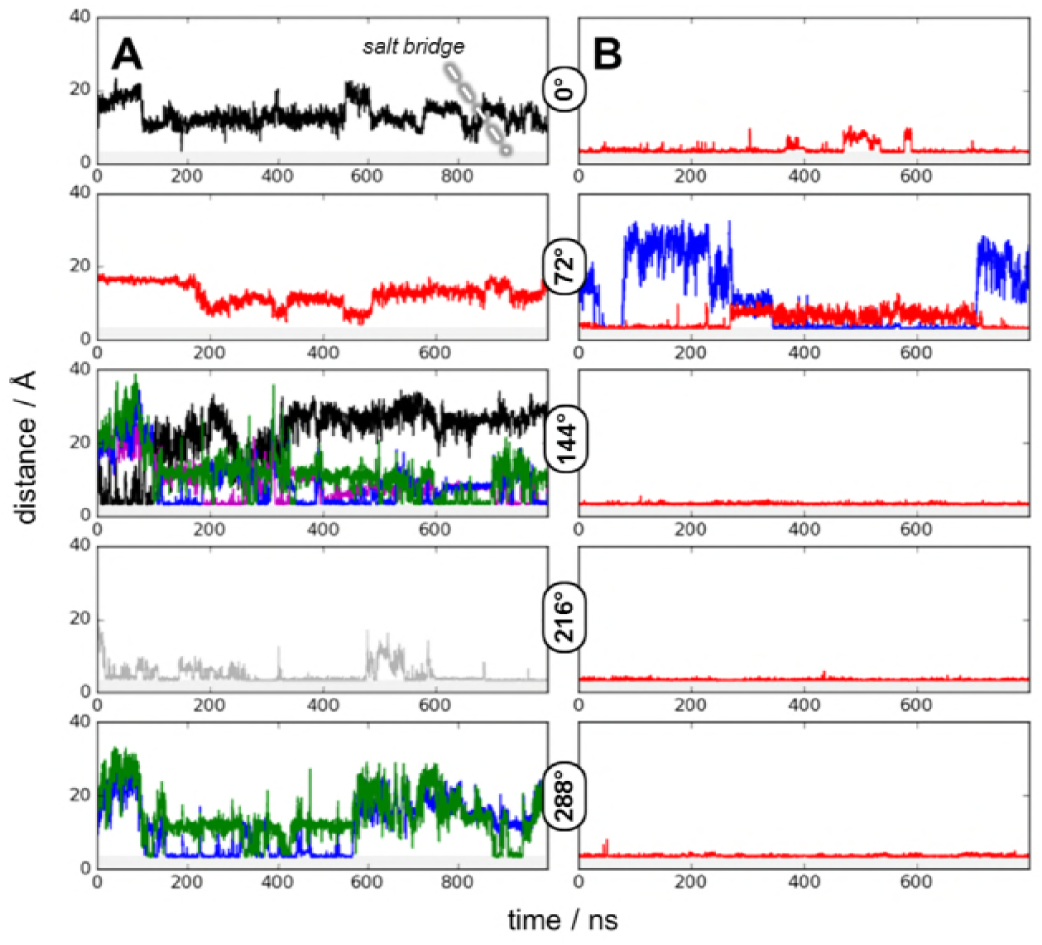
*i*+*3* salt bridge formation plays a key role in stabilization of secondary structures in pHLIP-1. A) Time-dependent salt bridge formation between R11 and sidechains of acidic residues in wt-pHLIP for 0°, 72°, 144°, 216°, and 288° orientations. For several orientations, formation of salt bridges between R11 and other residues is transient (0°, 72°, and 288°) or switches between multiple acidic residues (144°). Only the 216° orientation forms a stable salt bridge for hundreds of ns, but it is least likely to form secondary structures (Fig. 3). *Black*: E3; *red*: D14; *grey*: D25; *magenta*: D31; *blue*: D33; *green*: E34; *grey background*: minimum distance for salt bridge formation. B) Time-dependent salt bridge formation between R11 and sidechains of acidic residues in pHLIP-1 for 0°, 72°, 144°, 216°, and 288° orientations. Even though there are more acidic residues on the N-terminus of pHLIP-1 that could be salt bridge partners, the only salt bridge that is formed is between R11 and D14. Not surprisingly, this also corresponds to the region in pHLIP-1 that forms a helical turn when bound to POPC. *i*+*3* salt bridges like the R11-D14 salt bridge have been shown to stabilize *α*-helical peptides in solution (Baldwin PNAS 1982, Baldwin PNAS 1987, Garcia 2003 BJ), either through energetic stabilization of the backbone hydrogen bond needed to form a helical turn (Baldwin PNAS 1982) or exclusion of solvent from the backbone hydrogen bond by the proximity of the sidechains (Scheraga PNAS 2000). Color scheme is the same as in A.

Another key finding from the DUVRR study was that pHLIP undergoes desolvation in state II (12). We observe similar behavior through a general decrease in the solvent-accessible surface area (SASA) upon binding to the bilayer surface. Although wt-pHLIP has variable partitioning into the POPC bilayer based on the orientation of the bound complex (Fig. 2A), large portions of the peptide undergo dehydration, illustrated by a decrease in SASA. In the 0° and 72° orientations, the N-terminal segment of wt-pHLIP is most dehydrated, while in the 216° orientation the sinker stretch is most dehydrated, and in the 288° orientation, the majority of the peptide is dehydrated (Supporting Information Fig. S4A). Dehydration can be a hallmark of partitioning or folding: for example, the N-terminus of the 0° orientation is partitioned into the headgroup region, dehydrated, and folds into an *α*-helix. However, the N-terminal portion of the 288° orientation is not well-partitioned but is dehydrated and folded into a helical turn. In contrast, regardless of orientation, regions of dehydration of pHLIP-1 are consistent with areas that 1) partition into the POPC bilayer and 2) fold into a helical turn (i.e., the middle portion of the peptide) (Fig. S4B). Finally, the SASA of these peptides in comparison to their behavior is solution may also explain why pHLIP-1 undergoes folding more readily than wt-pHLIP: pHLIP-1 has a markedly lower SASA in state I, requiring less than a 30% reduction in SASA from state I to state II, whereas wt-pHLIP must undergo almost 50% reduction in SASA (Fig. S4).

### pHLIP utilizes multiple types of non-bonded interactions in state II

Further inspection shows that R11 plays multiple roles in both wt-pHLIP and pHLIP-1 yet is not a requirement for formation of a bound complex. It is possible for R11 to form hydrogen bonds or salt bridges with the phosphatidylcholine headgroup in POPC. For wt-pHLIP, R11 can form from three to six hydrogen bonds with the PC headgroups (Supporting Information, Figs. S5A and S6A). These hydrogen-bonded interactions may be enhanced by stable salt bridge formation; in particular, the 0° and 72° orientations have salt bridges that are stable for hundreds of ns. Not surprisingly, these orientations show the most stable binding to POPC (Figure 2). Similarly, in pHLIP-1 R11 can form multiple hydrogen bonds as well as stable salt bridges over hundreds of ns (Figs. S5B and S6B). This behavior correlates with stable binding of the surrounding residues in pHLIP-1 of each of the three orientations, again showing that R11 can play an important role in the formation of pHLIP-bilayer complexes (i.e., state II). Finally, other residues also contribute to hydrogen-bonded interactions of pHLIP with the bilayer; for example, pHLIP-1 forms multiple stable hydrogen bonds in the 0° and 72° orientations, none of which involve R11.

The transmembrane segment of wt-pHLIP and pHLIP-1 (residues 8-30 according to solid-state NMR experiments (14)) has a dual topology. In particular, the N-terminal portion has several aromatic sidechains (Y8, W9, Y12, and W15) that possess highly favorable partitioning free energies (36). Based on the depth of partitioning of W9 and W15, it appears that there is no discernible behavior with respect to the tryptophan sidechains that enhance binding to the POPC bilayer (Supporting Information, Fig. S7). However, a closer examination tells a different story. If we characterize the relationship between orientation of the tryptophan sidechain and the depth of partitioning into the POPC bilayer, W9 in wt-pHLIP adopts two distinct conformations (~40° and ~120°, slightly oriented towards and away from the bilayer normal) at the level of the phosphorus atoms in the PC headgroups (Fig. S8A). W15 in wt-pHLIP also adopts two major orientations, with the dominant population at 60° leading to favorable partitioning into the headgroups (~5 Å below the P atoms). In contrast, there is no apparent correlation between the tryptophan sidechain orientation and depth of binding for pHLIP-1 (Fig. S8B). Extending this analysis to Y8 and Y12 yields similar results: both Y8 and Y12 in wt-pHLIP have sidechain orientations that adopt a bimodal distribution. The populations that have sidechain orientations pointed inward and nearly parallel to the membrane normal (i.e., < 45°) also possess the most effective binding (Fig. S9A). Y8 in pHLIP-1 is similar to W9 and W15, with no preferred orientation and lying above the headgroup region (Fig. S9B). Interestingly, the dominant population of Y12 in pHLIP-1 occurs when the tyrosine sidechain adopts an orientation that is parallel to the bilayer surface; this orientation does not allow the sidechain to intercalate between lipids, unlike the other aromatic sidechains in either pHLIP peptide. This preferred orientation may be introducing a localized defect in the membrane surface, a potential insertion mechanism for several types of cell-penetrating peptides (37).

### Relating simulation results to theory

Wimley and White developed a scale for determining the free energy of partitioning from the aqueous phase into a POPC bilayer for amino acid residues (36). Aromatic and nonpolar amino acids have the most favorable partitioning free energies, while polar (and in particular, charged) residues have the most unfavorable partitioning free energies. Using this scale, we can predict the segment of wt-pHLIP and pHLIP-1 that will most favorably bind to the bilayer (Fig. 5). The most favorable stretch of wt-pHLIP that can partition into the bilayer is residues 8–31 (Δ*G*_partition_ = −6.6 kcal/mol, Table S1). This stretch corresponds remarkably well with the residues that partition most effectively into the POPC surface (see 0° orientation, Figure 2). Likewise, for pHLIP-1, the most favorable stretch that can partition into the bilayer is residues 8-30. Note that in pHLIP-1, the favorable stretch includes the C-terminus, most likely due to the absence of charged residues that are present in wt-pHLIP.

**Figure 5:**
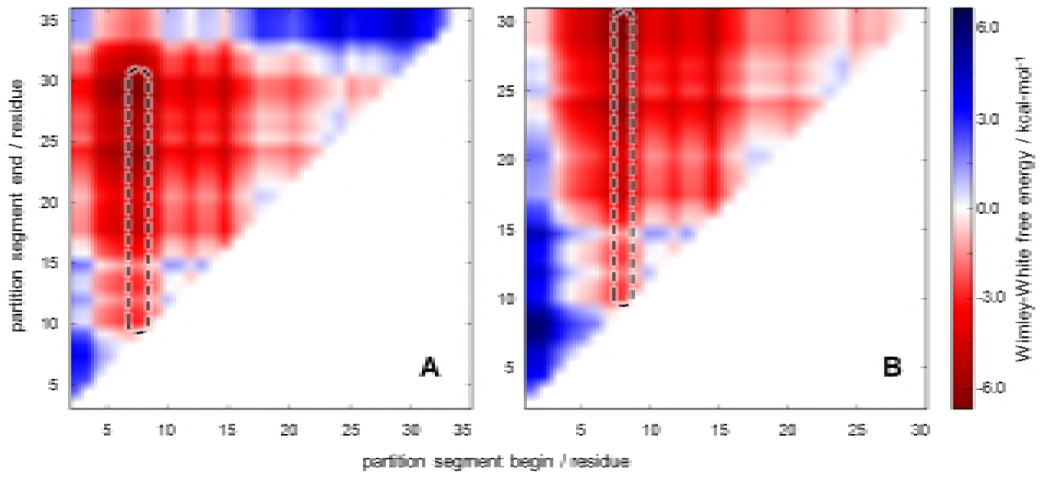
Wimley-White partition free energies provide insight into optimal binding of pHLIP. A) Theoretical per-residue cumulative partition free energy of wt-pHLIP. Values are based on the scale for free-energy partitioning of amino acid residues into membranes developed by Wimley and White (36). *x-axis*: starting point in wt-pHLIP for inclusion of per-residue partitioning free energies. *y-axis*: ending point in wt-pHLIP for inclusion of per-residue partitioning free energies. B) Theoretical per-residue cumulative partition free energy of pHLIP-1. Dashes indicate the segment of each peptide that possess the most favorable partitioning free energies.

Comparing the per-residue depth of partitioning for wt-pHLIP shows that in general, the C-terminal end of wt-pHLIP remains distal from the P atoms in the PC headgroups, with slight partitioning of the peptide from residues ~17-23 for the 0°, 216°, and 288° orientations (Figs. 2 and S1). The residues in wt-pHLIP with the most negative partitioning free energies (i.e., the aromatic residues Y8, W9, Y12, and W15) do not exert a localized partitioning effect. However, stretches of residues favorable to partitioning (e.g., the leucine residues from position 21 to 29) have a more proximal effect on the ability of wt-pHLIP to bind and partition into the POPC surface. Similar to the C-terminus of wt-pHLIP, the N-terminal half of pHLIP-1 predominantly lies away from the PC headgroups, due to the presence of acidic residues. However, this repulsion is compensated by partitioning of the middle and C-terminal portions of pHLIP-1 into the bilayer (Supporting Information, Fig. S1B). It appears that partitioning of the C-terminal “sinker stretch” is more pronounced in pHLIP-1 compared to wt-pHLIP, most likely due to the absence of acidic residues (D31, D33, and E34) on the C-terminus of pHLIP-1.

## DISCUSSION

In this study we have computationally modeled state II of wt-pHLIP and the variant, pHLIP-1. This is the first computational study of its kind, as previous MD studies have focused on insertion of *α*-helical pHLIP (38, 39), conformational sampling of gold nanoparticle-pHLIP conjugates (40), and pHLIP in solution (14, 19). In contrast, the field of antimicrobial peptides and cell-penetrating peptides has been extensively studied by MD simulations (41–43). This distinction of pHLIP from other CPPs is mainly tied to the anionic versus cationic nature of these two classes of peptides: the ability of acidic residues to quickly undergo titration to protonated states allows pHLIP to make rapid transitions from state II (coiled, bound) to state III (helical, inserted) when exposed to acidic conditions (18, 44). However, explicitly changing protonation states is not easily accomplished with conventional MD simulation techniques, since classical mechanics do not allow for formation and scission of the O–H bond in acidic amino acid residues. This shortcoming is also compounded by the long timescales associated with pHLIP insertion (ms–s), as even state-of-the-art membrane protein simulations are only capable of reaching tens to hundreds of *µ*s (45, 46). We have chosen to focus on the binding step in pHLIP function, as it avoids these two issues.

We can glean important details about the binding mechanism of pHLIP based on several structural studies. The first study, using brominated phospholipids, determined that the tryptophan residues of pHLIP (W9 and W15) are embedded at different depths within the bilayer (11). The DUVRR study of Cooley, Jiji, and coworkers determined that pHLIP undergoes partial helical formation in state II, most likely in conjunction with dehydration of the peptide backbone (12). In addition, the REDOR NMR study of An, Qiang, and coworkers focused on interactions of ^13^C-labeled nonpolar residues (A10, A13, L21, L22, L26, A27) with POPC (13). They found that the N-terminal residues (A10, A13) were in close proximity to the phosphatidylcholine headgroups in both state II and state III, whereas the C-terminus most likely remained unbound to the bilayer surface until the pH reached slightly acidic levels (pH 6.4). Although they observed two populations of conformations at pH 6.4, the C-terminus (i.e., the “sinker stretch”) was helical for both, indicating that helix formation most likely initially occurs around the D25 protonation switch. In their latest study, ^13^C solid-state NMR was used to determine that the C-terminal aspartic acid residues undergo titration first (i.e., highest p*K_a_*’s), leading An, Qiang, and coworkers to postulate that the solvent-exposed residues on the C-terminus underwent protonation under acidic conditions more quickly than residues in the middle of the peptide (D14 and D25) (14). By combining the insights gained from our simulations with these experimental observations of pHLIP, we can begin to formulate a general mechanism for binding of pHLIP in state II.

The first principle in understanding state II of pHLIP based on our simulations is that location of negatively-charged residues matters. In wt-pHLIP, the three negatively-charged residues on the C-terminus (D31, D33, and E34) undergo repulsive interactions with the PC headgroups; this relationship is the opposite for pHLIP-1, since there are negatively-charged residues on the N-terminus (E2, D3, and D6) (Fig. 2). These unfavorable interactions are conserved in all of our simulations. A second factor that contributes to effective partitioning of pHLIP is hydrophobic interactions. The nonpolar residues from L21 to V30 form the “sinker stretch” posited by An and coworkers (13) and is bisected by D25. Using the hydrophobicity scale proposed by Wimley and White (36), the sinker stretch has a favorable cumulative free energy of partitioning from water to the membrane of −2.45 kcal-mol^−1^ (Supporting Information, Table S1). If we include the C-terminal acidic residues present in wt-pHLIP, the free energy of partitioning becomes unfavorable (+2.03 kcal-mol^−1^). Clearly, the ability of the sinker stretch to partition into the bilayer can be influenced by the number of anionic residues present in the C-terminus of pHLIP. However, the presence of these residues do not lead to complete dissociation of wt-pHLIP, as the sinker stretch remains bound to the POPC surface in the majority of our simulations (Fig. 2A).

Other non-bonded interactions play a critical role in partitioning of pHLIP in state II, specifically the N-terminal half of the transmembrane segment. The presence of several aromatic residues (Y8, W9, Y12, and W15) and the lone cationic residue (R11) near the D14 protonation switch lead to a noticeably different binding mechanism from the sinker stretch in pHLIP. (Interestingly, the N-terminal half of the transmembrane segment in wt-pHLIP and pHLIP-1 has a more favorable free energy of partitioning (−3.67 kcal-mol^−1^) than the sinker stretch (Supporting Information Table S1).) In addition to the aforementioned DUVRR study showing significant desolvation of wt-pHLIP in state II, the fluorescence spectra of pHLIP-1 in state II were more intense than wt-pHLIP, indicating that the tryptophans of pHLIP-1 become more deeply embedded within the bilayer upon binding (12, 18). In our simulations, tryptophans play different roles for wt-pHLIP and pHLIP-1: in wt-pHLIP, one or both tryptophan residues partition below the surface and discarding solvent molecules only when the sidechain is parallel to the membrane normal, while in pHLIP-1 the orientations of tryptophans appear to have little effect on binding. Similar behavior is seen with the tyrosine residues, with Y12 in pHLIP-1 playing a potential role in creating a local defect in the bilayer surface. Binding of aromatic sidechains is favored due to a combination of factors (the quadrupolar moment of the indole ring, the hydrophobic effect, and the complex electrostatic environment of the headgroup region (47)). In lieu of binding through interactions of aromatic sidechains with the bilayer, pHLIP-1 appears to utilize R11 to form a salt bridge with D14 to facilitate partitioning. R11 may be further stabilizing partitioning by forming salt bridges with the PC headgroups; this is a common mechanism for stabilizing small peptide-membrane interactions (e.g., the Kras protein (48)). The combination of nonbonded interactions with the bilayer illustrates how a peptide like pHLIP can partition into the membrane surface, even in the presence of charged residues.

Finally, binding of pHLIP to the membrane is enhanced by formation of helical turns. Partitioning-folding coupling for cationic peptides is well-characterized, with a reduction of −0.4 kcal-mol^−1^ per residue for a peptide to undergo folding into a helix while bound to a bilayer surface (49). In a similar way, the folding that occurs in both pHLIP peptides contributes to favorable partitioning into the bilayer. Our observation of stable helical turns in pHLIP-1 is consistent with the original investigation of pHLIP-1, which detected the presence of *α*-helical folds in CD spectra of state II (18). What was unexpected is that we observed formation of stable helical turns in wt-pHLIP in several of our simulations as well (Fig. 3A). The fact that wt-pHLIP and pHLIP-1 undergo folding into helical turns in different segments of the peptide is consistent with proposed models of the reversible state II to state III transition (14, 39, 50, 51). In wt-pHLIP, folding mainly occurs from I7-Y12 and D14-T18 but also in the sinker stretch from L21-V30. In pHLIP-1, folding is almost completely conserved from Y12-T18 and can extend from L22-V30. Based on solid-state NMR and fluorescence spectroscopy studies, the C-terminal half of the transmembrane segment of pHLIP is most sensitive to changes in pH ((14, 51) and can form a peripheral helix prior to insertion (50). In addition, recent constant-pH MD studies that were initiated in state III showed that at pH 7, the N-terminal half of the transmembrane segment (i.e., including D14) diffuses towards the surface of the bilayer but remains in an *α*-helical fold(39). Regardless of the location of folding (i.e., centered around D14 or D25), it appears that wt-pHLIP and pHLIP-1 are undergoing a priming action for the transition to state III: helical formation can occur while either proton switch is deprotonated and the hydrophilic side chain is solvent-exposed. In the case of D25, upon acidification and protonation of the residue (a transition from +1.23 to −0.07 kcal-mol^−1^ in partitioning free energy), the C-terminus of pHLIP becomes continuously nonpolar, facilitating the more extensive helical formation that is the hallmark of the transition from state II to state III. Protonation of D14 would lead to a similar outcome, as that portion of the transmembrane segment also possesses a favorable partitioning free energy. The propensity for each of these regions to undergo partial helical folding, even in state II, is not surprising; our recent study showed that pHLIP spends a noticeable portion of time sampling helical conformations in solution (19). It is conceivable that pHLIP could be partially folded when binding to a membrane surface.

Multiple pathways exist for pHLIP to bind to the cell membrane surface, but preferential binding is governed by specific biophysical characteristics of the peptide (Fig. 6). Before the transition to acidic pH and state III, the sinker stretch of the C-terminus of the peptide begins to bury itself beneath the headgroup region and into the hydrophobic interior of the bilayer. In our simulations, this effect can be modulated by the presence (wt-pHLIP) or absence (pHLIP-1) of acidic residues. Furthermore, it is clear that binding via the N-terminal half of pHLIP can also occur. This interaction may require more extensive sampling of potential peptide-bilayer configurations, since binding takes place through a combination of hydrophobic effects (aromatic side chains) and electrostatic ones (R11). Our results indicate that state II is the result of a two-step binding process, one in which both halves of the transmembrane helical domain (bisected by P20) undergo burial into the bilayer interface. A more general issue we have begun to address is the complexity of how pHLIP interacts with a cell membrane. The field has progressed from the conventional three-state model to begin to account for differences in membrane composition from POPC (15–17) as well as utilizing advanced spectroscopic techniques that allow for a more detailed characterization of pHLIP conformational sampling in the bound and inserted states (13, 14, 50, 51). A combination of experimental and computational approaches will be required to fully understand how pHLIP functions, which will subsequently enable design of more specific and effective variants of pHLIP.

**Figure 6:**
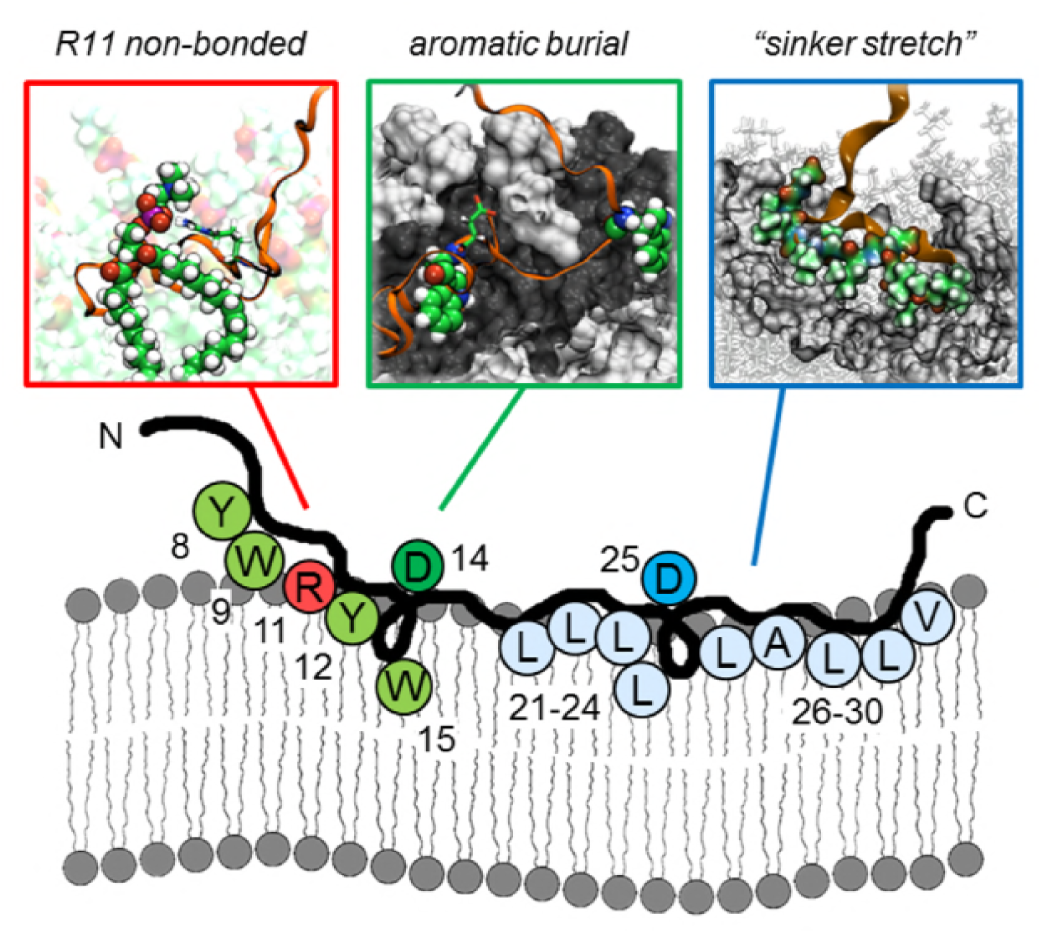
Proposed binding mechanism of pHLIP to POPC. pHLIP utilizes a combination of non-bonded forces to achieve binding to the cell membrane surface. *Blue*: the C-terminal half of pHLIP uses its “sinker stretch” of hydrophobic residues to embed below the level of the lipid headgroups, sequestering itself from aqueous solvent. This interaction is responsible for the helical priming action observed around the D25 protonation switch. *Inset*: colored spheres represent sinker stretch residues that interact with the headgroups and acyl chains of the POPC bilayer upper leaflet, grey surface. *Green*: burial of the aromatic sidechains of Y8, W9, Y12, and W15 within the PC headgroups helps anchor the middle portion of pHLIP to the bilayer surface. These interactions stabilize pHLIP around the D14 protonation switch and allow for helical formation. *Inset*: interaction of W9/W15 (spheres) with the headgroup region of the POPC bilayer. *Red*: hydrogen bonding or salt bridge formation of R11 with the PC headgroup can work in conjunction with aromatic burial to stabilize binding of pHLIP. *Inset*: R11 (sticks) forming a hydrogen bond with the phosphate of a PC headgroup (spheres).

## CONCLUSION

The computational studies carried out here provide strong support for the experimental biophysical studies characterizing state II of pHLIP. Intimate knowledge at the molecular level of detail is fundamental to identifying specific interactions that govern the free energy release upon binding of pHLIP to the cell membrane surface. This free energy release forms one component of the thermodynamic cycle associated with the “partitioning-folding coupling” process first proposed by Wimley and White (36). They postulated that with this fundamental understanding, it would be possible to formulate a governing set of rules for the rational design of peptides with specific partition coefficients and secondary structures(52). Efforts are currently underway in our lab to extend computational investigations of this thermodynamic cycle and bring Wimley and White’s goal to fruition.

## AUTHOR CONTRIBUTIONS

CG, YR, and BM designed the research. CG and YR carried out all the simulations and analyzed the data. CG, YR, and BM wrote the article.

## ACKNOWLEDGMENTS

The authors would like to thank Austin Clark for critical reading of the manuscript. This work was supported by West Virginia University (B.M.) and NIH MCB-1714888 (C.G., B.M.). Computational time was provided through WVU Research Computing and XSEDE allocation no. TG-MCB 130040.

## SUPPLEMENTARY MATERIAL

An online supplement to this article can be found by visiting BJ Online at http://www.biophysj.org.

